# Impact of Ferumoxytol Magnetic Resonance Imaging on the Rhesus Macaque Maternal-Fetal Interface

**DOI:** 10.1101/699835

**Authors:** Sydney M. Nguyen, Gregory J. Wiepz, Michele Schotzko, Heather A. Simmons, Andres Mejia, Kai D. Ludwig, Ante Zhu, Kevin Brunner, Diego Hernando, Scott B. Reeder, Oliver Wieben, Kevin Johnson, Dinesh Shah, Thaddeus G. Golos

## Abstract

Ferumoxytol is a superparamagnetic iron oxide nanoparticle (SPION) used off-label as an intravascular magnetic resonance imaging (MRI) contrast agent. Additionally, ferumoxytol-uptake by macrophages facilitates detection of inflammatory sites by MRI through ferumoxytol-induced image contrast changes. Therefore, ferumoxytol-enhanced MRI holds great potential for assessing vascular function and inflammatory response, critical to determine placental health in pregnancy. This study sought to assess the fetoplacental unit and selected maternal tissues, pregnancy outcomes, and fetal well-being after ferumoxytol administration. In initial developmental studies, pregnant rhesus macaques were imaged with and without ferumoxytol administration. Pregnancies went to term with vaginal delivery and infants showed normal growth rates compared to control animals born the same year that did not undergo MRI. To determine the impact of ferumoxytol on the maternal-fetal interface, fetal well-being, and pregnancy outcome, four pregnant rhesus macaques at ∼100 gd (gestational day) underwent MRI before and after ferumoxytol administration. Collection of the fetoplacental unit and selected maternal tissues was performed 3-4 days following ferumoxytol administration. A control group that did not receive ferumoxytol or MRI was used for comparison. Iron levels in fetal and maternal-fetal interface tissues did not vary between groups. There was no significant difference in tissue histopathology with or without exposure to ferumoxytol, and no effect on placental hormone secretion. Together, these results suggest that the use of ferumoxytol and MRI in pregnant rhesus macaques will not introduce a detectable risk to the mother or fetus at the time of imaging or up to one year following normal vaginal delivery.

**Summary Sentence:** Ferumoxytol magnetic resonance imaging for non-invasive pregnancy monitoring of the rhesus macaque does not impact histopathology or iron content of the maternal-fetal interface.

## Introduction

In the hemochorial human and nonhuman primate placenta, maternal intervillous blood bathes the placental villi, allowing oxygen and nutrient transfer to the fetal blood circulating within the capillaries of the villous stroma. Pregnancy complications may stem from maladaptation of maternal vessels causing insufficient placental perfusion, leading to macrophage recruitment, cytokine release, and hypoxia at the maternal-fetal interface (MFI). Compromised intervillous flow is associated with adverse pregnancy outcomes [1-4]: insufficient placental perfusion could result in fetal growth restriction, preeclampsia, and pregnancy loss. The ability to identify abnormal uteroplacental vascular adaptation, compromised perfusion, and attendant inflammation could be valuable in identifying at-risk pregnancies before clinical manifestations.

Currently, ultrasound is the most commonly used method to assess fetal growth. It is also used to detect umbilical and uteroplacental blood flow abnormalities, but only indirectly through velocity waveform analysis. Further, ultrasound lacks the ability to detect immune cell homing to the MFI that may precede adverse pregnancy outcomes. Magnetic resonance imaging (MRI) can provide high-resolution anatomic and functional information including blood velocities and flow, perfusion, and oxygenation to characterize placental implantation site, visualize maternal pelvic structures, and diagnose abnormally aggressive trophoblast invasion or placental abruption [5]. In many clinical applications, gadolinium-based contrast agents (GBCAs) are used to quantify tissue perfusion and perform high-resolution angiography [6]. In the non-human primate, gadolinium MRI has been used to investigate spiral artery and perfusion domain (cotyledon) location, and quantify placental perfusion [7,8], the latter of which has also been achieved in humans [9]. However, GBCAs have been shown to cross the placenta into the fetus with uncertainty in the long-term consequences of in utero GBCA exposure. Although there is no specific evidence that it causes teratogenic or chromosomal damage [5,10-12], the risk to the fetus of gadolinium based MR contrast agent administration remains unknown and should not be routinely provided to pregnant patients [13].

We explored an alternative approach for quantitative tissue perfusion and MR angiography in pregnancy, using the SPION ferumoxytol as a contrast agent. Ferumoxytol is approved for the treatment of iron deficiency in adults, including pregnant women. It has also emerged as an off-label MR contrast agent with favorable MR properties [14,15] that can yield high-detail angiography and functional information about the MFI non-invasively, including quantitative perfusion maps of maternal blood that allow for analysis of individual cotyledons in the placenta [16], as seen in imaging with gadolinium [7]. As such, it has high potential to identify local and global perfusion abnormalities that might be indicative of placenta pathologies..Our initial MR imaging results suggest that ferumoxytol stays within the maternal blood and does not cross the placenta into the fetal circulation immediately after ferumoxytol administration [17]. Ferumoxytol also has the potential to spatially localize inflammatory events, as the nanoparticles are taken up by activated cells of the mononuclear phagocyte system at sites of tissue inflammation, which can then be imaged after ferumoxytol in the blood space has cleared [14,18-22]. The MRI transverse relaxation rate R2* has a known linear relationship with the concentration of iron in tissues. Therefore, R2* mapping may enable localization of iron-laden macrophages, as well as quantification of their density.

Ferumoxytol has been previously used in, but is not limited to, the study of inflammation of the pancreas in patients with type-1 diabetes [22], inflammation of the lymph nodes in patients with Hodgkin lymphoma [18,21], inflammation in patients with osteomyelitis and arthritis, the study of normal adrenal function, monitoring of kidney transplant vessel patency, monitoring intracranial aneurysms for potential rupture, and in the tracking of stem cell grafts. Application of ferumoxytol use to the MFI may be extremely valuable for the monitoring of placental dysfunction. Ferumoxytol is routinely used to treat anemia in pregnant mothers. Its safety profile and properties for MR imaging makes it a promising contrast agent to fill a gap in the non-invasive diagnosis of placental health with potential for clinical routine use. Importantly, demonstrating the safe use of ferumoxytol for placental imaging is a necessary step in this application. Therefore, the purpose of this work is to assess the feasibility of ferumoxytol administration on the MFI, fetal well-being, and pregnancy outcomes in a non-human primate model. The rhesus macaque provides an accurate experimental model of the human MFI and immune system, having hemochorial placentation, endovascular trophoblast invasion with attendant spiral artery remodeling, and chorionic villous placental architecture. We observed no negative impact from ferumoxytol injection on the histopathology at the MFI, or evidence of ferumoxytol transfer to the fetus, as assessed by iron content in MFI and fetal tissues These results demonstrate the feasible infusion of ferumoxytol in a cohort of pregnant rhesus macaques. Future work will utilize this instructive animal model with ferumoxytol-enhanced MRI to challenge experimental paradigms and understand interventions assessing placental function and pregnancy well-being.

## Materials/Methods

Several aspects of the impact of ferumoxytol on the Rhesus Macaque MFI were interrogated: in vitro analysis of immune cell isolation and incubation with ferumoxytol; placental explant incubation with ferumoxytol; maternal and fetal outcomes at year post-birth for rhesus that underwent ferumoxytol MRI during pregnancy vs. controls; tissue iron content, maternal plasma analysis, and histopathology analysis in a cohort of rhesus that went fetectomy after undergoing ferumoxytol MRI.

### Immune Cell Isolation and Incubation with Ferumoxytol

In vitro ferumoxytol-uptake studies using monocytes and macrophages were isolated from whole blood drawn from pregnant rhesus macaques, at approximately 100gd, as previously reported [23]. Neutrophils were isolated as previously published [24]. All three cell types were incubated in ferumoxytol (Feraheme, AMAG Pharmaceuticals, Waltham, MA) at 0, 50, 100 or 200 µg/ml for 1 hour. Additionally, there were incubations of 0 µg/ml or 200 µg/ml with activating agents (50 ng/ml phorbol-12-myristate-13-acetate (PMA; Sigma-Aldrich, St. Louis MO)) for all cell types, 750 ng/ml ionomycin (Sigma-Aldrich, St. Louis MO) for monocytes only). Following incubation, cells were washed and fixed with 2% paraformaldehyde (PFA) for visualization of iron content by Prussian Blue staining [25-27]. Cells were imaged using a Nikon Eclipse TE300 microscope with NIS-Elements image capture.

### Placental Explant Incubations in Ferumoxytol

Prior to imaging experiments, placental explants were prepared from tissues obtained from untreated animals undergoing fetectomy or caesarean section in unrelated studies, during first trimester of pregnancy or at term. Explants were incubated in ferumoxytol at 0, 100 or 200 µg/ml, diluted in DMEM/F12 with 10% fetal calf serum, for 2, 4, and 24 hours at 37°C in room air/5% CO2. Explants were fixed in 2% PFA overnight and embedded in paraffin blocks. Tissues were imaged using a Nikon Eclipse TE300 microscope with NIS-Elements image capture.

### Prussian Blue Staining

To visualize cellular iron content, isolated, fixed immune cells grown on coverslips or deparaffinized rehydrated tissue sections were incubated in Prussian blue solution [25-27] for 20 minutes, washed with deionized water, and mounted with Aquapolymount (Polysciences, Warrington PA).

### Care and Use of Macaques

Female rhesus macaques in the Wisconsin National Primate Center (WNPRC) breeding colony were housed with compatible males and monitored for breeding and menses. Pregnancies were confirmed and dated (+/− 2 days) based on menstrual cycle, observation of copulation, and ultrasound measurements of gestational sac and fetuses. Blood samples were collected using a needle and syringe or vacutainer system from the femoral or saphenous vein. All macaques were cared for by WNPRC staff in accordance with the regulations and guidelines outlined in the Animal Welfare Act and the Guide for the Care and Use of Laboratory Animals. This study was approved by the University of Wisconsin-Madison Graduate School Institutional Animal Care and Use Committee (IACUC).

### MRI Impact on Pregnancy Outcome and Postnatal Growth

There were two imaging phases in this study. In the first phase (Supplemental Fig. 1 A), seven pregnant macaques were imaged, three with ferumoxytol and four without, to establish standard methods for anesthesia, imaging, and pilot scan settings with pregnancies proceeding to term. Infants joined the WNPRC colony and body weights during the first year of life were compared to 116 untreated macaque infants born at the WNPRC in 2016 to determine if MRI during pregnancy impacted postnatal growth. Mean and standard deviation for these colony infants were calculated at different ages, similar to the approach previously published [28]. Weights of infants exposed to MRI with or without ferumoxytol in-utero that were then born into the colony were compared to this WNPRC 2016 colony growth chart through their first year of life.

### Use of IL-1b to Induce MFI Inflammation

In the second phase (Supplemental Fig. 1 B), we used a paradigm of ferumoxytol MRI following intra-amniotic injection of 10mg IL-1β (has been reported previously to increase decidual macrophage numbers and model chorioamnionitis and preterm labor [29,30]) or sterile saline (n=4) to test the efficacy of ferumoxytol detection of mononuclear phagocytes and inflammation [14,18-22] at the MFI. Untreated controls (n=4) did not receive intra-amniotic injections, ferumoxytol, or MRI. While the resulting R2* maps from IL-1β-exposed animals did not differ from those of animals receiving intra-amniotic saline (Zhu et al, under review), comparison of saline-injected and untreated animals allows determination of any impact of ferumoxytol MRI at the MFI or on the fetus.

### Intra-amniotic Injection

Procedures were performed under transabdominal ultrasound guidance on the lateral aspect of the abdomen. A syringe filled with sterile saline was attached to a biopsy needle and inserted through an aseptically prepared site of the abdominal wall until the tip reached the wall of the uterus, avoiding the bowel and bladder (n=4, Supplemental Fig. 1 B). The needle was advanced into the amniotic cavity and a small amount of amniotic fluid was drawn to confirm needle placement. The contents of the syringe were then slowly injected into the amniotic cavity, and the needle was withdrawn. Following withdrawal of the needle, the insertion site in the uterus was observed by ultrasound to confirm lack of bleeding.

### MRI

All animals that underwent an MRI exam, regardless of whether they also received ferumoxytol, were sedated by injection of up to 10 mg/kg ketamine, intubated, and anesthesia was maintained by inhalation of oxygen and 1.5% isoflurane. A pulse oximeter probe was placed and vital signs were monitored every 15 minutes. Animals were imaged in the right-lateral position and a respiratory bellow was placed around the animal’s belly during imaging to enable respiratory-compensated imaging that minimizes motion-related artifacts. Animals that received ferumoxytol had an intravenous catheter placed for injection during imaging.

Animals that received ferumoxytol, dynamic contrast enhanced (DCE) images were acquired on a clinical 3.0T MRI system (Discovery MR750, GE Healthcare, Waukesha, WI) utilizing a 32-channel torso radiofrequency coil (Neocoil, Pewaukee, WI). Time resolved T1-weighted DCE images with 5 second temporal resolution were obtained throughout the ferumoxytol administration [17]. Ferumoxytol diluted 5:1 with normal saline was administered at 4 mg/kg body weight over a 20 second interval using a power injector, followed by a 20 ml saline flush at the same rate. A baseline R2* MRI scan (an MRI relaxation parameter highly correlated and sensitive to detect iron concentration) was performed before ferumoxytol administration. R2* measurements were estimated in the maternal, MFI, and fetal tissues by region-of-interest analysis directly from the MRI images. Follow-up R2* MRI scans were performed on subsequent days after contrast injection to determine the persistence of ferumoxytol in various tissues. MRI acquisition parameter details can be found elsewhere [17 *and* Zhu et al, under review].

### Fetectomy

At ∼gd100 the fetoplacental unit was collected via hysterotomy (n=8, Supplemental Fig. 1 B). Maternal biopsies were collected aseptically during surgery and the dam recovered. The fetus was euthanized by intravenous or intracardiac injection of 50 mg/kg sodium pentobarbital. Fetal and MFI tissues were dissected for histology, iron content mass spectrometry, and protein assay.

### Tissue Homogenates

Tissues collected at fetectomy (0.1-0.7g) (n=8, Supplemental Fig. 1 B) were homogenized in a Bullet Blender (Next Advance, Troy NY) at full power for 10 minutes with non-metal blending beads and 500 µL PBS. Tissue homogenate was stored at −80°C until use. A 96-well format micro BCA protein assay (Thermo Scientific, 23235) was used to determine protein concentrations in homogenates assayed for iron content according to the manufacturer’s instructions.

### Iron content determinations

Tissue homogenates were assayed for iron concentrations at the Wisconsin State Laboratory of Hygiene Trace Element Research Group in selected maternal, fetal, and MFI tissues via inductively coupled plasma - optical emission spectrometry [31-34]. The limit of detection is 1μg/g tissue.

### Steroid Hormone Extraction and LC/MS/MS Analysis

Maternal plasma samples (450 μl) collected for multi-steroid analysis from animals that had tissues collected at fetectomy (Supplemental Fig. 1 B) were extracted and assayed as previously reported [35,36].The limit of detection is 30pg/ml for progesterone; 6pg/ml for estrone and estradiol.

### Histology

Tissues collected for histology were fixed in 4% PFA overnight, 70% ethanol overnight, and routinely processed and embedded in paraffin. 5µm sections were stained with H&E and assessed by veterinary pathologists blinded to treatment groups. Tissues were evaluated for the presence or absence of physiologically significant pathologic changes, normal anatomic variations, and inflammation. Morphologic diagnoses (Supplemental Data 2) summarize these histologic findings. Organs not given a morphologic diagnosis are considered to have no significant pathologic or inflammatory changes and were scored as a 0. Severity (none=0, minimal=1, mild=2, moderate=3, severe=4) was determined by the extent and distribution of inflammation, vascular change (infarction, thrombosis, pregnancy associated vascular remodelling and/or the lack thereof), and non-vascular necrosis across the tissue section or organ (multiple slides were necessary to evaluate the placenta). Scores were averaged and compared between treatment groups as previously reported [37]. Some MFI tissue sections were stained with Prussian Blue for iron localization.

### Statistics

Iron concentrations in tissue homogenates were compared between treatment groups by 2-way ANOVA and Sidak’s multiple comparison test. Differences in pathology and changes in R2* values were assessed by 2-way ANOVA. Hormone level changes were assessed by 1-way ANOVA.

## Results

### PBMC Incubations

To determine whether rhesus macaque cells take up ferumoxytol as reported with human cells [14,18-22], prior to initiating the imaging phases of this study, rhesus monocytes, macrophages, and neutrophils were incubated in 100 µg/ml ferumoxytol (Fig. 1), the approximate concentration of ferumoxytol in the blood with administration for MRI, and iron was visualized by Prussian blue staining. Staining was seen in differentiated macrophages but not monocytes or neutrophils. Activation with PMA and ionomycin did not affect iron staining. No staining was seen without ferumoxytol incubation.

**Fig. 1.**
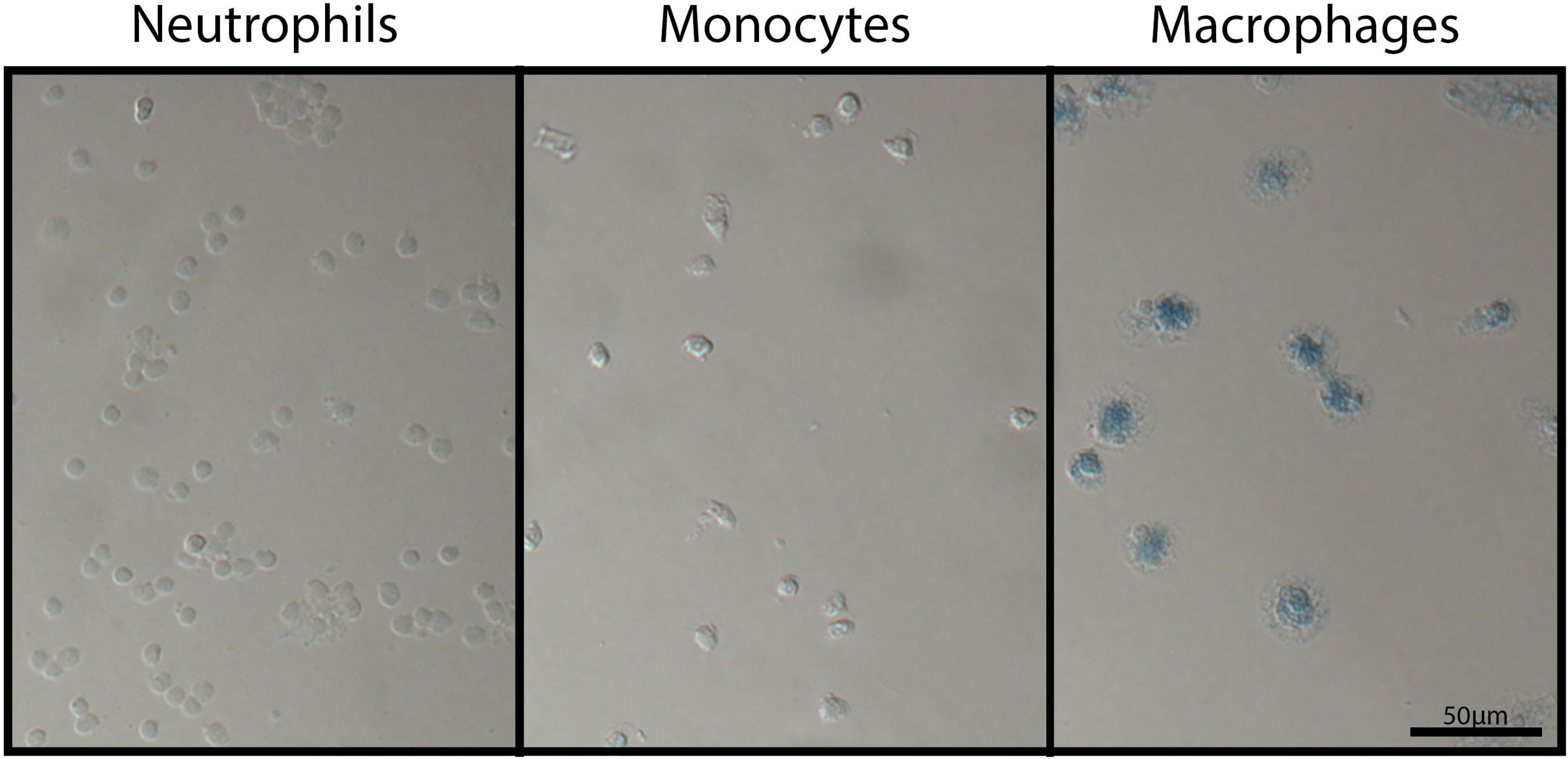
Localization of iron in rhesus immune cells incubated with ferumoxytol. Peripheral blood neutrophils (left), monocytes (center), and *in vitro*-differentiated macrophages (right) from rhesus macaque whole blood were incubated in 100 µg/ml ferumoxytol for 1 hour and stained with Prussian Blue. Photomicrographs are of cytospins of neutrophils, or monocytes or macrophages grown on coverslips in culture.

### Placental Explants Incubations

To determine whether placental ferumoxytol uptake by placental tissue may confound use for inflammation mapping in vivo, prior to initiating the imaging phases of this study, rhesus placental explants were incubated with ferumoxytol and stained with Prussian Blue. Modest background Prussian Blue iron staining in tissue was observed independent of ferumoxytol incubation, likely indicating endogenous iron content (Supplemental Fig. 2). Minimal increase in iron staining was observed after 2 hours of ferumoxytol-incubation. An increase in iron staining appeared after 24 hours incubation, specifically in the villous endothelium of the placental tissue. Not substantial staining of the syncytiotrophoblasts was observed, the primary interface exposed to ferumoxytol in maternal blood in vivo. Low levels of endogenous iron and modest increases in ferumoxytol uptake in control placental explants ex vivo after incubations suggests that in vivo inflammation detection by ferumoxytol-enhanced MRI would be feasible and not confounded by background placental iron content/uptake.

### Maternal Clinical Outcomes with Ferumoxytol Administration

In addition to the 7 animals in this study that received ferumoxytol for MRI (Supplemental Fig. 1), 28 pregnant rhesus monkeys from other ongoing studies (unpublished) had up to three ferumoxytol imaging sessions. In 35 total experimental subjects who had ferumoxytol imaging sessions, two animals required moderate medical attention following ferumoxytol administration. Both animals had periocular edema following IV bolus administration of ferumoxytol that was treated with 10 mg diphenhydramine hydrochloride. One animal had a short period of increased heart rate and SPO2 levels. This animal had previous ocular swelling not associated with ferumoxytol, so it is unclear whether this event was due to ferumoxytol or other drugs used to anesthetize the animal. These mild allergic reactions responded to diphenhydramine and the animals recovered without further medical intervention.

### Pregnancy Outcomes

Seven pregnant rhesus macaques (Supplemental Fig. 1 A) who underwent MRI gave birth via vaginal delivery at term, and the infants joined the WNPRC colony. Results of these imaging studies are described in separate reports [17,38]. Pregnancy outcomes were generally unremarkable, with one retained placenta (which occurs in ∼2.6% of WNPRC pregnancies). None of the seven dams had immediate or long-term reactions to the ferumoxytol treatment.

Infant growth data from these pregnancies are plotted along with their birth year cohort weights (Fig. 2). The weights of the MRI offspring generally stayed within one standard deviation of the average infant weights. Infants followed normal physiological and sociobehavioral patterns seen in other colony infants as assessed by daily veterinary observations.

**Fig. 2.**
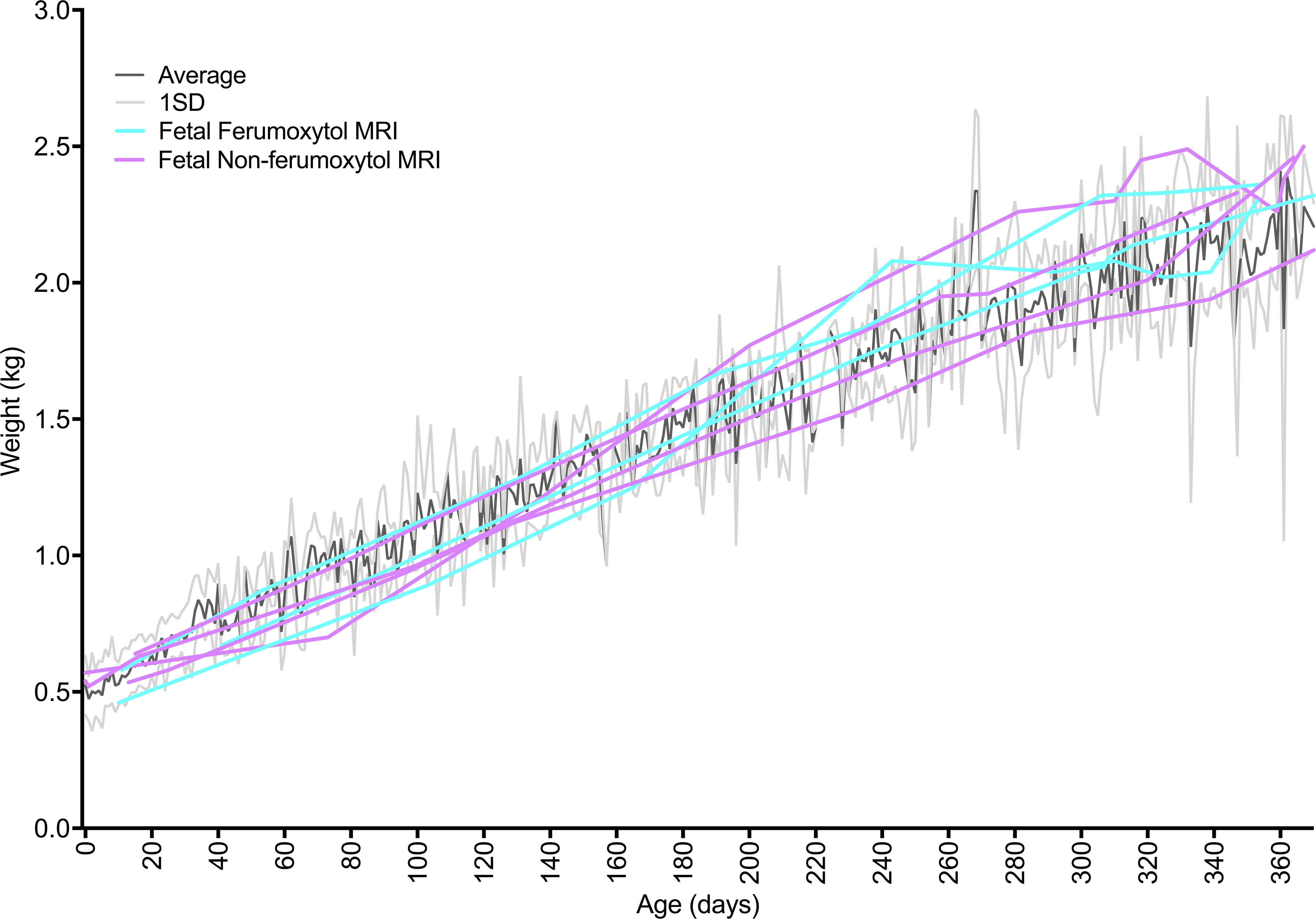
Infant growth rates with maternal ferumoxytol treatment compared to animal colony controls. The black line represents the mean weight (kg) for 116 infants born at the WNPRC in 2016, weighed at the age in days listed on the x-axis. The grey lines represent one standard deviation from the mean. Purple lines represent one animal each that was imaged by MRI without ferumoxytol. Aqua lines represent one animal each that was imaged with ferumoxytol, plus 4 additional scans without additional ferumoxytol administration. The irregular mean and standard deviation lines reflect the fact that not all colony animals were weighted on any given day so the data represent a different population of animals at any specific time point.

### Ferumoxytol Detection by MRI Following Administration

Three of the seven pregnant rhesus macaques that carried infants to term (Supplemental Fig. 1 A) had been imaged with ferumoxytol at ∼100gd. Imaging occurred immediately before (to establish a baseline R2* values in maternal, MFI, and fetal tissue) and 15 minutes after administration of ferumoxytol, followed by four follow-up MRI scans at approximately one day, one week, two weeks and three weeks following ferumoxytol administration. In all three animals, an increase in R2* values in both the primary and secondary placental disks is seen immediately following ferumoxytol injection (Fig. 3). he R2* values in fetal lung remained close to baseline though all scans while fetal liver R2* values increased slightly in two of the three animals. This may reflect an increase in physiological iron transport to the fetus over time in normal pregnancy, unrelated to ferumoxytol. The R2* values in the placenta, which increased dramatically following ferumoxytol administration, returned to approximate baseline levels within one day post-ferumoxytol, supporting a rapid clearance of ferumoxytol from the blood. Ferumoxytol accumulation in the placenta or transfer to the fetus was not detectable by R2* (Zhu et al, under review).

**Fig. 3.**
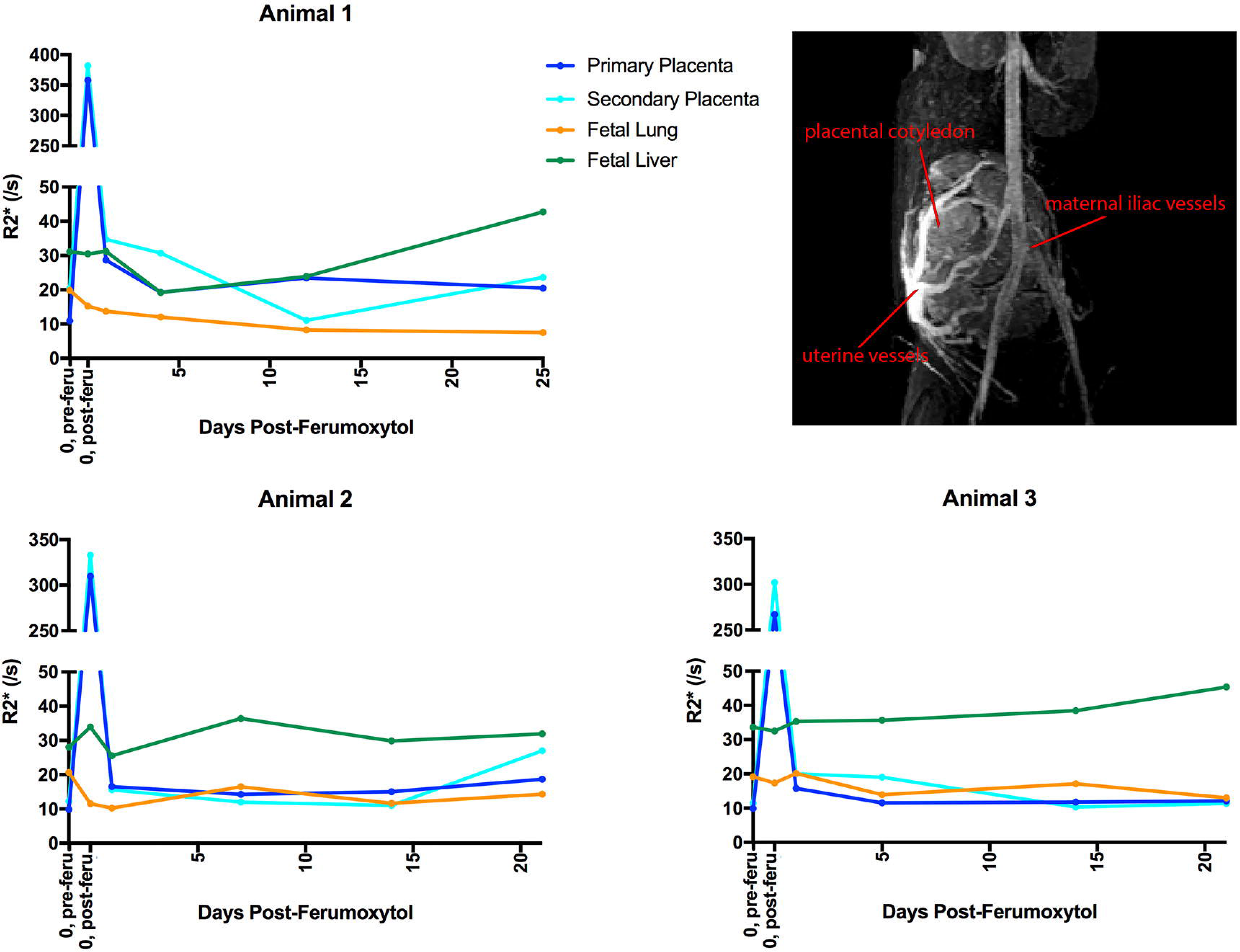
R2* values following ferumoxytol injection. R2* values were monitored in 3 pregnant rhesus macaques immediately following and 1 day, 1 week, 2 weeks, and 3 weeks ferumoxytol injection. The image on the top right is a representative Dynamic Contrast Enhanced (DCE) image of maternal and uterine ferumoxytol detection, including placental intervillous flow, illustrating the imaging data used to determine R2* values. The first point represents pre-injection (“0, pre-feru”) and the second point represents the same day post-injection (“0, post-feru”). Primary placental disc values are in dark blue, secondary placenta in aqua, fetal lung in orange, and fetal liver in green.

### Iron Content in Tissues

Maternal, MFI, and fetal tissues from 8 pregnancies (Supplemental Fig. 1 B) were surgically collected at ∼gd100 following MRI and iron concentrations were determined in these tissues (Fig. 4). When ferumoxytol-exposed and untreated control groups were compared, only maternal liver showed a significant increase in iron concentration with maternal ferumoxytol administration over control (*p*<0.0001). There are no significant differences in iron in fetal tissues in ferumoxytol vs. non-ferumoxytol-exposed animals.

**Fig. 4.**
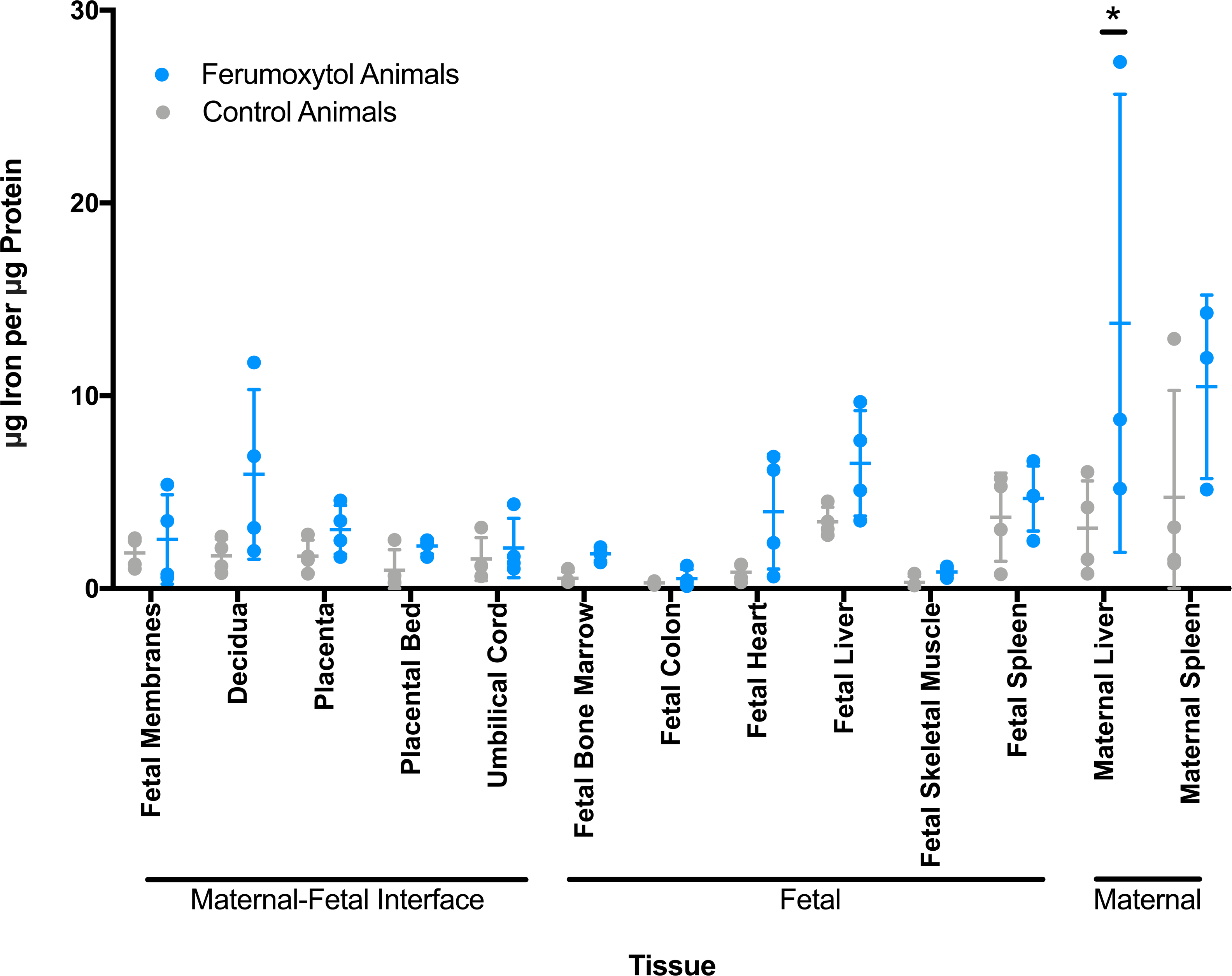
Iron content of maternal and fetal tissues. Iron content of selected tissues was determined by mass spectrometry. Non-imaged animals are represented by grey circles (n=4). Animals that received ferumoxytol imaging with intra-amniotic saline are in blue (n=4, except for maternal liver and maternal spleen where n=3). Mean and standard error are denoted by horizontal lines for each tissue.

### Prussian Blue Staining of MFI Tissues

Prussian Blue staining varied animal-to-animal in the placenta, decidua, and fetal membranes from animals that underwent fetectomy. Tissues from ferumoxytol-receiving animals, overall, did not have noticeably different staining compared to non-ferumoxytol-receiving animals. Interestingly, the animal with the most consistent staining had not received ferumoxytol (Supplemental Fig. 3), likely reflecting normal physiological iron.

### Ferumoxytol Effects on Plasma Progesterone, Estrone, and Estradiol

For each imaging day in animals that underwent fetectomy (Supplemental Fig. 1 B), maternal plasma samples were assessed by mass spectrometry for progesterone, estrone, and estradiol levels [35,36] to assess the impact of MRI imaging and ferumoxytol administration on placental endocrine function. Non-imaged controls (Supplemental Fig. 1 B) received a one-time plasma-collection at time of fetectomy. There was no statistically significant change in placental hormone levels following administration of ferumoxytol and hormone levels generally stayed within the range of levels seen in non-imaged controls (Fig. 5).

**Fig. 5.**
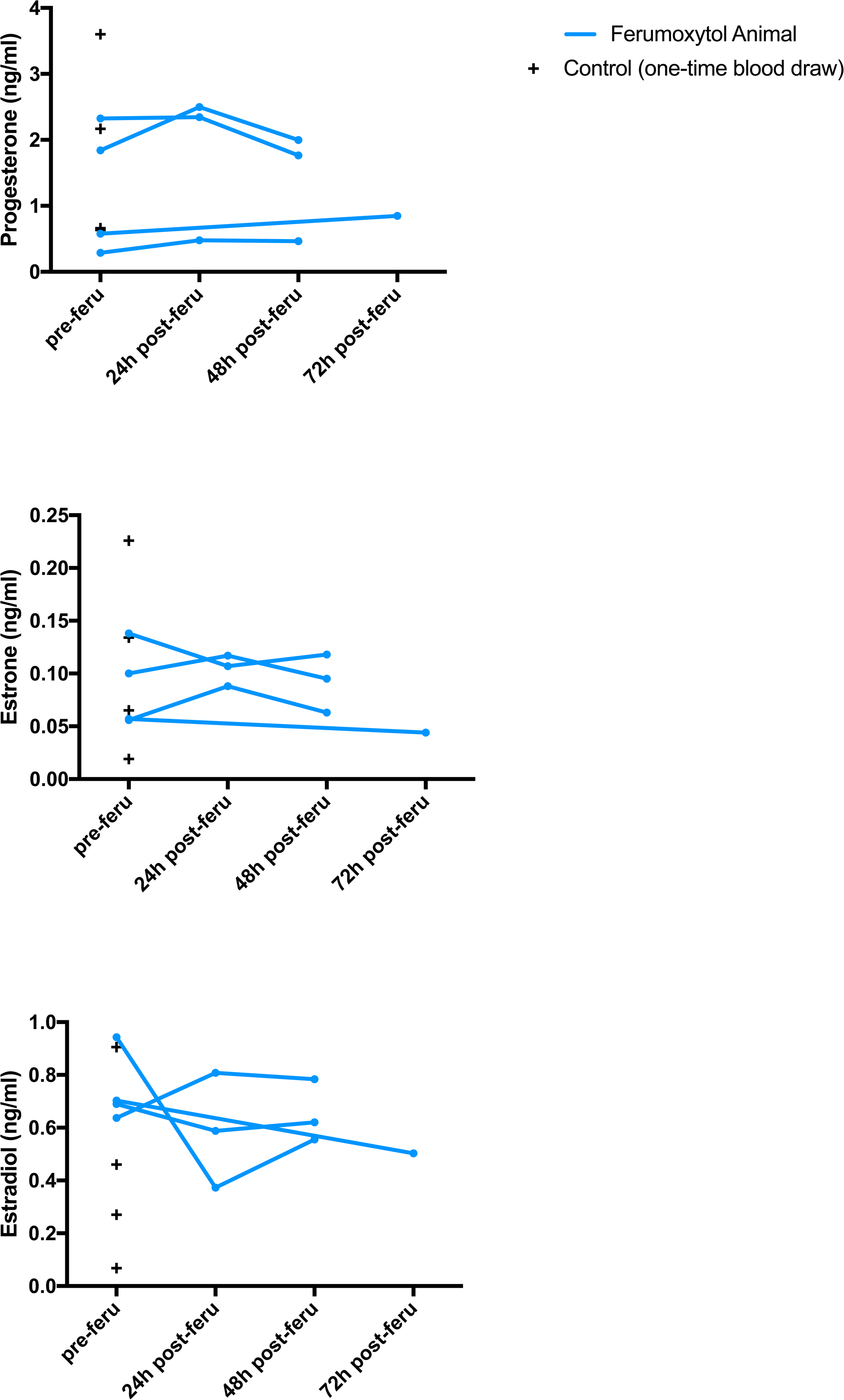
Plasma hormone levels in MRI animals assessed by mass spectrometry. Blue lines represent progesterone, estrogen, and estradiol levels in ferumoxytol-infused animals before injection, 24h following injection, and 48h or 72h following injection. Black plus signs (+) represent single blood draw readings from non-ferumoxytol control animals, indicating the expected range of peripheral blood steroid hormone levels in pregnant macaques.

### Ferumoxytol Effects on Histopathology

Of 37 maternal and fetal tissues collected at fetectomy (Supplemental Data 1), the placenta, decidua, amniotic membranes, placental bed, maternal spleen, and maternal liver had notable histopathology. Animals that did and did not receive ferumoxytol MRI had no statistically significant differences in individual tissue histopathology scores (Fig. 6). Morphologic Diagnoses are provided in Supplemental Data 2.

**Fig. 6.**
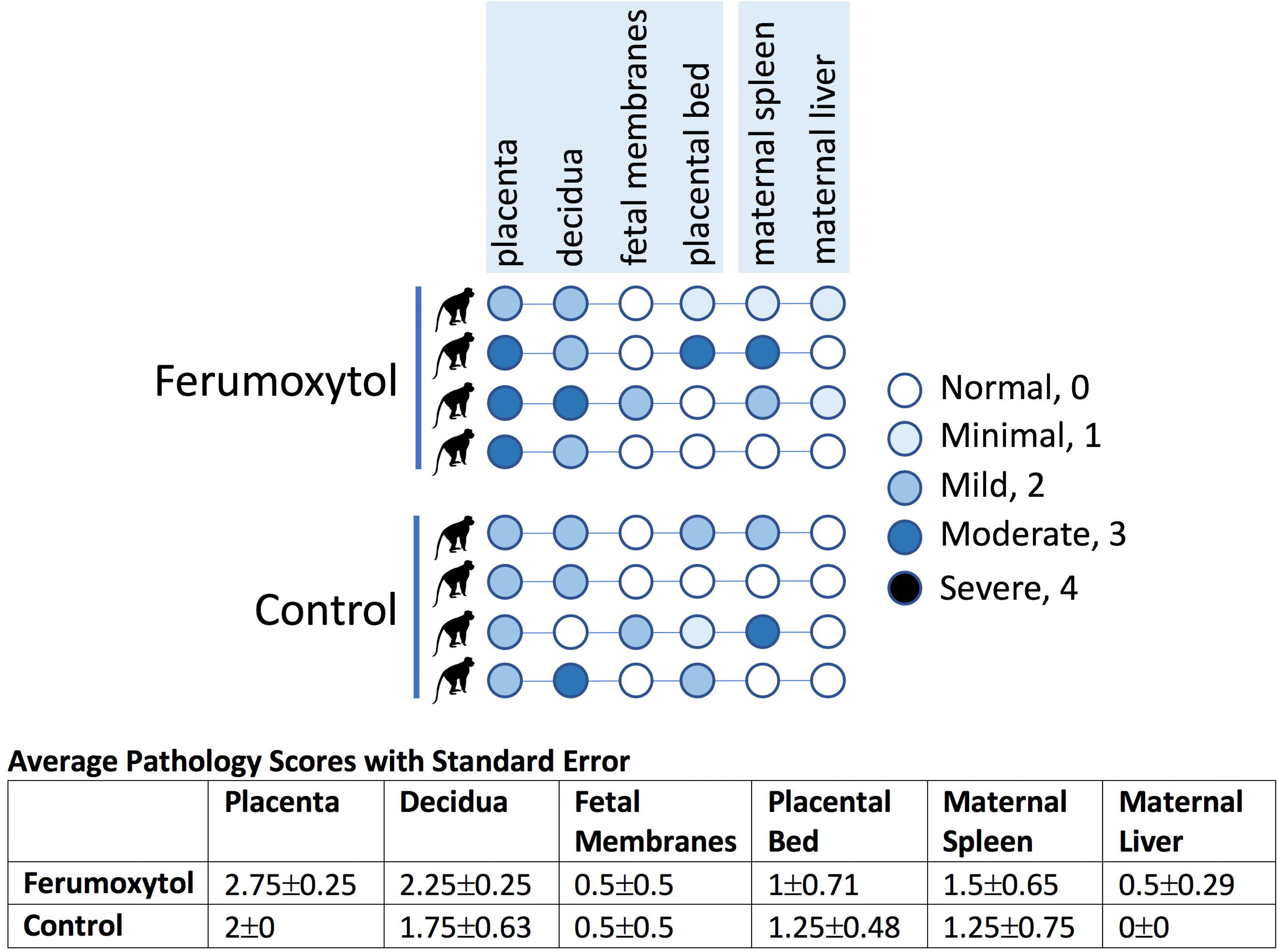
Chart summarizing histopathology scores of all tissues that showed pathology. For each animal, the circle representing each tissue is colored to denote severity of pathology. The top 4 rows represent the ferumoxytol-receiving intra-amniotic saline animals and the bottom 4 rows are for non-ferumoxytol non-MRI controls. Numbers were assigned to each severity rating and used to analyze pathologies (normal=0, minimal=1, mild=2, moderate=3, severe=4). The chart below presents the average pathology scores for each tissue per treatment, used to assess statistical significance.

## Discussion

In this study, we examined the impact of MRI with and without ferumoxytol on the fetoplacental and maternal tissues, pregnancy outcomes, and fetal well-being in the pregnant rhesus macaque. Offspring from imaged pregnancies with or without ferumoxytol had uneventful labor and normal growth in comparison with contemporary pregnancies from the WNPRC breeding colony. No significant impact of MRI with ferumoxytol on iron content or histopathology of fetal and MFI tissues (decidua, placenta, fetal membranes) was observed. Placental function as indicated by peripheral blood steroid hormone levels was unaffected by MRI with ferumoxytol. It should be noted that ferumoxytol was injected as a diluted bolus in this study, while it is administered as a slow infusion in humans to reduce the risk of anaphylactic reactions. The lack of significant adverse outcomes in the rhesus subjects also suggests the utility of ferumoxytol as a non-gadolinium contrast agent for MRI in pregnancy studies.

The use of ferumoxytol in MRI of pregnancy has the potential to yield important diagnostic information. Additionally, ferumoxytol-enhanced MR angiography allows for visualization of maternal uteroplacental vessels involved in transporting blood to and from the uterus, and therefore the placenta. When using ferumoxytol as the contrast agent for DCE imaging, the time of arrival of ferumoxytol-laden blood into the intervillous space and perfusion rates of blood into the individual cotyledons can be determined, which may be related to the health of the placental tissue [7,8]. Fetal vessels are not enhanced since significant ferumoxytol does not pass into the fetal circulation during MRI [17], observations further supported by fetal tissue iron data reported here.

We have shown that ferumoxytol is taken up by rhesus monkey phagocytic cells, demonstrating the feasibility of designing additional studies for its application to nonhuman primate models of adverse pregnancy outcomes. Ferumoxytol has not been applied previously to the nonhuman primate model. *In vitro* culture experiments demonstrated that the SPION was taken up by macrophages differentiated from peripheral blood monocytes, but not by undifferentiated monocytes or granulocytes. This indicates that ferumoxytol is a feasible reagent to detect the accumulation of phagocytic macrophages at sites of inflammation. Tissue macrophages take up ferumoxytol and clear these iron nanoparticles more slowly than those in the blood, therefore, sites of inflammation can be located by performing delayed imaging following ferumoxytol administration. This paradigm may help identify inflammation at the MFI, which could predict an insult to the pregnancy.

Concerns about ferumoxytol uptake by the placenta and the potential for transport of elevated levels of iron to the fetus, putting the fetus at risk for hemochromatosis or pulmonary hemosiderosis since these disorders result in fetal growth restriction, hepatic failure, alveolar hemorrhage, and stillbirth [39-41] were addressed. Placental villous explants were incubated *in vitro* with physiologically realistic concentrations of ferumoxytol, and staining of explant tissue sections for iron content with Prussian Blue did not demonstrate any significant uptake of SPION by placental tissues in a physiologically meaningful pattern (i.e., syncytiotrophoblast uptake) that would be anticipated with exposure of the placenta to ferumoxytol in the maternal blood in the intervillous space. With *in vivo* treatment of pregnant rhesus macaques, there was no significant impact on maternal health, pregnancy outcome, or postnatal fetal development. Pilot studies used to establish the parameters for imaging, indicated that offspring from survival pregnancies showed uneventful labor and normal fetal/infant growth compared to contemporary pregnancies from the WNPRC breeding colony. Furthermore, the dose of ferumoxytol used in these animal studies, while allowing sensitive imaging of the MFI, is quite low (4 mg/kg) compared to human therapeutic dosing for anemia. This underscores the expected safety of ferumoxytol in this pregnancy model.

The fetus acquires iron during pregnancy through transferrin receptor acquisition of ferritin and transit across the placental syncytiotrophoblast and cytotrophoblast to the fetal vasculature within villous stroma [42]. Placental tissues collected from MRI experiments and stained with Prussian Blue for iron content did not reveal discernible differences between tissues from control and ferumoxytol-treated pregnancies. Additionally, decidual tissues and fetal membranes did not demonstrate any consistent differences between experimental groups. There were focally distributed areas of iron detected by Prussian Blue staining, however interestingly, the tissues with the clearest demonstration of iron content were the decidua and fetal membranes rather than the placental villi. It is important to note that the animal in which iron was most readily demonstrated in these tissues did *not* receive ferumoxytol and thus SPION-delivered iron was not the source of Prussian Blue staining. These data suggest that although the placenta directly transports iron to the fetus via a biologically conserved ferritin/ferritin receptor-mediated pathway, this active pathway does not participate in the uptake of ferumoxytol by the syncytiotrophoblasts. While the mechanism of ferumoxytol’s uptake by macrophages has not been determined, similar dextran-coated SPIONs are taken up by phagocytosis or SR-A-mediated endocytosis [43-45]. We hypothesize that cellular iron sequestration, as indicated by Prussian Blue staining, may be largely attributable to macrophage uptake of erythrocytes as a routine surveillance function at the MFI.

Consistent with a lack of increase in iron content of MFI tissues by histochemical methods, there was no significant increase in iron concentration in MFI tissues by mass spectrometry. Likewise, fetal tissues that would be anticipated to accumulate iron, did not show a statistically significant increase. While there does appear to be a trend for slightly higher, though not statistically significant, iron content in fetal tissues, further studies will be needed to determine if this is a consistent result. There was a statistically significant increase in maternal liver iron content, which was expected since the liver is a main clearance organ for ferumoxytol, with resident hepatic macrophages (Kupffer cells) taking up ferumoxytol particles in studies in rabbit [46] and human subjects [47,48].

Histopathology was evaluated in selected maternal tissues, the MFI, and in fetal tissues. There was no detectable histopathology in any fetal tissues. While histopathology was noted in tissues at the MFI, there were not significant differences between ferumoxytol-receiving and control animals. Some histopathological features were noted among placentas even in untreated “normal” pregnancies. This lack of a difference in pathological findings in placental, decidual, and fetal membrane specimens from the animals in study also supports the use of ferumoxytol in this animal model. Functional assessment of the placenta by monitoring of placental hormone secretion (progesterone, estradiol, estrone) likewise revealed no significant difference between animals receiving ferumoxytol MRI imaging, and untreated animals.

The data presented in this report were part of a larger study in which some fetuses received IL-1B via an intra-amniotic injection with the goal of inciting trafficking of inflammatory phagocytes to the MFI [29,30]. Ferumoxytol MRI and histochemical and mass spectrometry analyses did not support an increase in iron-retaining cells at the MFI. This previously published model reported increased numbers of macrophages and granulocytes in the decidua parietalis, however that study did not evaluate the decidua basalis which we hypothesized would be imaged with Ferumoxytol treatment. While our study did not validate the use of Ferumoxytol with this model due to lack of induced inflammation, it is possible that other nonhuman primate models of adverse pregnancy outcomes and MFI inflammation, including maternal infection with *Listeria monocytogenes* [49] or Zika virus [37,50-52] which have been shown to provoke significant inflammation in the decidua basalis with significant placental pathology, may more productively demonstrate the efficacy of ferumoxytol for detection of inflammation at the MFI. Furthermore, a placenta facing a bacterial or viral insult may have altered placental transporter protein expression, which may affect ferumoxytol’s ability to pass into the fetal blood circulation [53]. Use of these pregnancy models in ferumoxytol MRI may reveal important experimental utility of the SPION.

The use of an animal model to evaluate MRI methodologies has significant advantages. The pregnant dam is anesthetized for the imaging procedure in the nonhuman primate model, and the inhaled anesthetic is transferred to the fetus, which is also anesthetized. Therefore, the fetal motion is minimized in MRI of the animals, leading to reliable MRI results. This experiment setup provided unique opportunity to validate the feasibility of MRI methodologies without the fetal motion being a confounding factor. However, anesthesia is not the standard of care for MRI evaluation of pregnant humans. The potential fetal motion in MRI of pregnant human subjects need to be addressed. A motion-robust R2* mapping technique has been proposed by our group and is under separate study (Zhu et al, under review). Upon successful validation, the motion-robust MRI technique may enable assessing detection macrophage homing in pregnant women. Other common MRI imaging strategies for motion include motion prevention (e.g. coaching, breath holding), imaging artifact reduction (e.g. physiological triggering and gating, fast imaging readouts), and motion correction (e.g. navigators, prospective/retrospective corrections) [54]. These, and other strategies, are commonly used in body imaging applications (i.e. cardiac, lung, abdominal) where motion is of substantial concern for producing diagnostic quality MR images.

In summary, we conclude that ferumoxytol administration for imaging in this rhesus pregnancy model is feasible. Future studies will explore the use of ferumoxytol to detect placental inflammation and the diagnostic value of DCE MRI in the presence of placental dysfunction. The rhesus macaque will be an important platform for initial development of novel imaging approaches in an experimentally tractable model.

## Acknowledgment

The authors wish to thank GE Healthcare who provides research support to the University of Wisconsin-Madison and AMAG Pharmaceuticals for providing Ferumoxytol for this study. Further, Dr. Reeder is a Romnes Faculty Fellow, and has received an award provided by the University of Wisconsin-Madison Office of the Vice Chancellor for Research and Graduate Education with funding from the Wisconsin Alumni Research Foundation.

## Supplemental Data Legends

**Supplemental Fig. 1. Experimental Design.** Monkey outlines represent a single animal that received each treatment, represented on their respective timeline. (A) Blue timelines outline experimental design for animals where pregnancy proceeded to term and the infants were born by spontaneous vaginal delivery. (B) Pink timelines outline the series of procedures that animals received whose pregnancies were terminated by fetectomy. IA=intra-amniotic, FTX=fetectomy.

**Supplemental Fig. 2. Histological analysis of rhesus placental explants.** Placental explants from tissue from first (left column) and third (right column) trimester pregnancies were incubated in ferumoxytol for 2 and 24 hours (200 µg/ml ferumoxytol for first trimester, 100 µg/ml ferumoxytol for third trimester). Original experiments were at the 200 µg/ml concentration but changed to 100 µg/ml as this concentration better reflects the concentration of ferumoxytol in the blood when imaging. Tissue explants were embedded in paraffin and sections were cut and stained with Prussian Blue to localize iron. The top row shows control tissue that was not incubated in ferumoxytol. The middle row shows the 2 hour incubation. The bottom row shows the 24 hour incubation.

**Supplemental Fig. 3. Histochemical analysis of iron at the MFI.** The left image presents a representative chorioamniotic membrane sample stained with Prussian Blue, the right image presents a full-thickness placental section (with decidua and membranes attached) similarly stained. Both samples were collected from a non-MRI, non-ferumoxytol control animal. The lower panels present a higher magnification view of the regions depicted by rectangles in the upper panels. These images are from an animal that did not receive ferumoxytol, the degree of Prussian Blue staining was not seen to increase in animals that received ferumoxytol (not shown).

**Supplemental Data 1. Tissues Collected at Fetectomy.** The table presents the tissues collected from dams and fetuses at fetectomy after ferumoxytol MRI, or from untreated pregnancies.

**Supplemental Data 2. Pathology Reports.** The table describes full pathology reports for the tissues summarized in Fig. 6.

