## Supplemental Data 1 for "Impact of Ferumoxytol Magnetic Resonance Imaging on the Rhesus Macaque Maternal-Fetal Interface"

**Supplemental Data 1: Tissues Collected at Fetectomy**

| **Maternal Biopsies** | Spleen, Liver, Mesenteric Lymph Node |
| --- | --- |
| **Maternal-Fetal Interface** | Placenta, Decidua, Placental Bed, Umbilical Cord, Chorioamniotic Membranes, Amniotic Fluid |
| **Fetal Central Nervous System** | Cerebrospinal Fluid, Frontal Cortex, Mid Cortex, Occipital Lobe, Cerebellum, Brain Stem, Cervical Spinal Cord |
| **Fetal Ocular** | Eye |
| **Fetal Cardiopulmonary** | Lung, Heart |
| **Fetal Reproductive** | Uterus or Prostate/Seminal Vesicles, Ovary or Testis |
| **Fetal Musculoskeletal** | Bone Marrow, Skeletal Muscle |
| **Fetal Immune** | Spleen, Thymus, Mesenteric Lymph Node, Tracheobronchial Lymph Node |
| **Fetal Gastrointestinal** | Liver, Duodenum/Jejunum, Ileum, Cecum, Colon |
| **Fetal Urinary** | Kidney |
| **Fetal Endocrine** | Adrenal, Pancreas, Pituitary |
