## Supplemental Data 2 for "Impact of Ferumoxytol Magnetic Resonance Imaging on the Rhesus Macaque Maternal-Fetal Interface"

**Supplemental Data 2: Pathology Reports**

|  | **Morphologic Diagnoses** |
| --- | --- |
| **Ferumoxytol 1** | - Placenta: mild multifocal basal plate thrombosis with mild acute multifocal villitis and intervillositis with increased intervillous fibrin, syncytial knots, moderate multifocal subchorionic fibrin and minimal segmental neutrophilic subchorionitis - Decidua: mild multifocal to diffuse acute (neutrophilic) deciduitis with minimal multifocal necrosis and hemosiderophages - Placental Bed: minimal multifocal lymphocytic endometritis - Spleen, maternal: minimal diffuse neutrophilic splenitis - Liver, maternal: minimal multifocal centrilobular lymphocytic hepatitis |
| **Ferumoxytol 2** | - Placenta: moderate multifocal basal plate thrombosis with minimal to mild multifocal neutrophilic intervillositis and villositis, with increased intervillous fibrin and syncytial knots and mild multifocal segmental subchorionic fibrin - Decidua: mild multifocal lymphocytic deciduitis with mild diffuse hemosiderophage accumulation - Placental Bed: moderate diffuse necrosuppurative superficial endometritis and mild multifocal lymphocytic perivascular myometritis - Spleen, maternal: moderate multifocal lymphoid hyperplasia |
| **Ferumoxytol 3** | - Placenta: moderate multifocal basal plate thrombosis, mild multifocal subchorionic fibrin, minimal multifocal acute subchorionitis, rare avascular villi, mild multifocal neutrophilic villitis and intervillositis - Decidua: moderate chronic lymphoplasmacytic deciduitis - Fetal membranes, decidua: mild multifocal chronic lymphoplasmacytic deciduitis with occasional neutrophils - Spleen maternal: mild diffuse neutrophilic splenitis - Liver, maternal: minimal periportal lymphoplasmacytic hepatitis with rare neutrophils |
| **Ferumoxytol 4** | - Placenta: moderate multifocal decidual arteriopathy with multifocal vascular fibrinoid necrosis and intraluminal thrombi with perivascular edema - Decidua: mild multifocal perivascular and interstitial lymphoplasmacytic deciduitis |
| **Non-ferumoxytol Control 1** | - Placenta: mild random multifocal villous avascularization, minimal multifocal mineralization, three small transmural infarctions - Decidua: mild multifocal lymphocytic deciduitis with rare neutrophils and focal occlusive fibrin thrombi - Placental Bed: mild multifocal lymphocytic endometritis - Spleen, maternal: mild diffuse neutrophilic splenitis |
| **Non-ferumoxytol Control 2** | - Placenta: mild retroplacental hemorrhage, mild multifocal basal plate infarction and vascular thrombosis with minimal multifocal mineralization and moderate basal plate hemorrhage - Decidua: mild multifocal chronic lymphoplasmacytic deciduitis |
| **Non-ferumoxytol Control 3** | - Placenta: mild multifocal remote (chronic) thrombosis and infarction with minimal acute (neutrophilic) inflammation and moderate multifocal mineralization - Fetal membranes: mild diffuse chorioamniotic hemosiderosis - Placental Bed: minimal neutrophilic vasculitis - Spleen, maternal: moderate multifocal lymphoid hyperplasia and mild diffuse neutrophilic splenitis |
| **Non-ferumoxytol Control 4** | - Placenta: mild neutrophilic intervillositis and villitis with syncytial knot formation; mild multifocal basal plate necrosis, hemorrhage, and mineralization; mild multifocal chorionic necrosis with acute inflammation - Decidua: moderate chronic lymphoplasmacytic deciduitis with marked organized hemorrhage - Placental Bed: mild multifocal vasculitis and perivasculitis |
