## Supplementary figures and images for "Impact of Ferumoxytol Magnetic Resonance Imaging on the Rhesus Macaque Maternal-Fetal Interface"

### Supplemental Figure 1

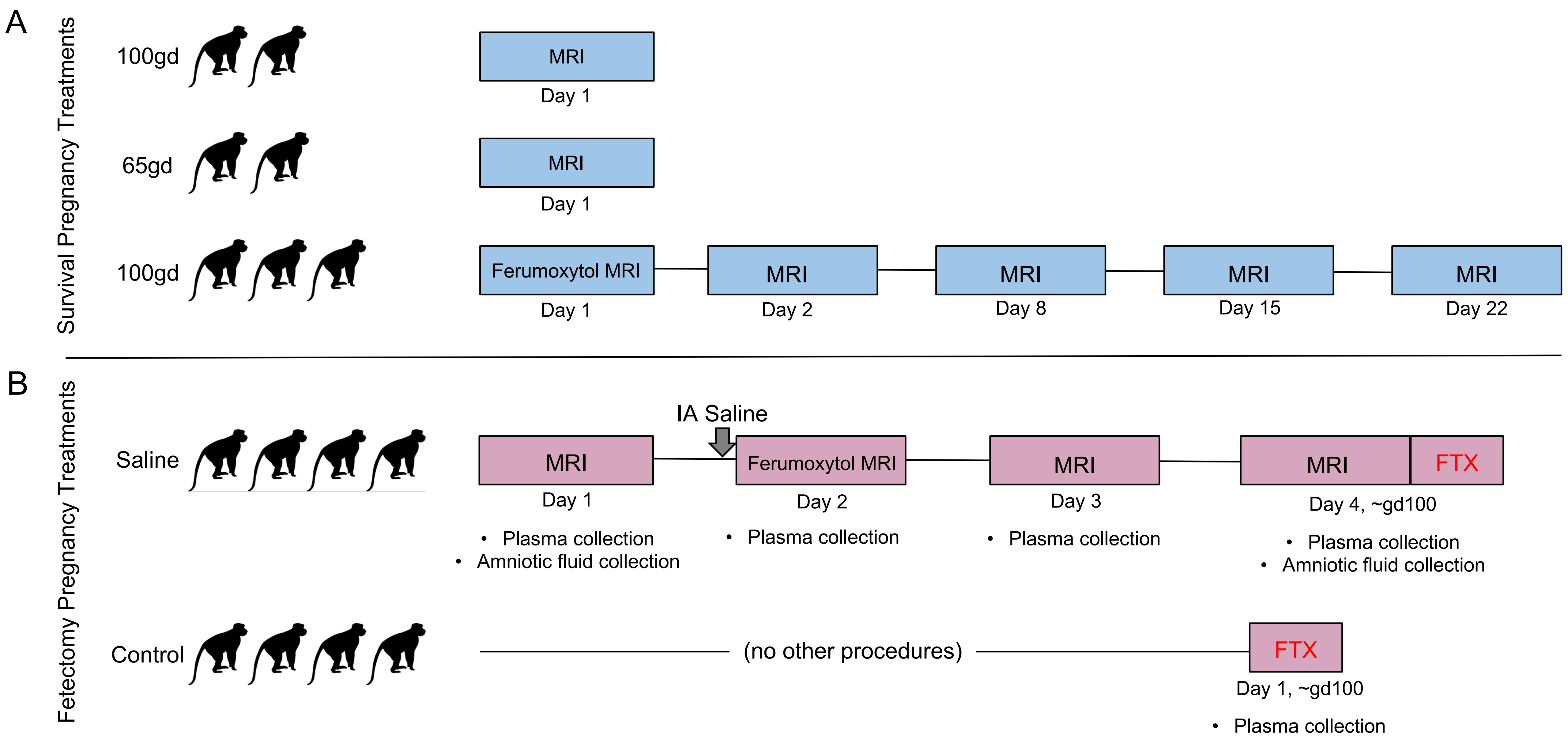

### Supplemental Figure 2

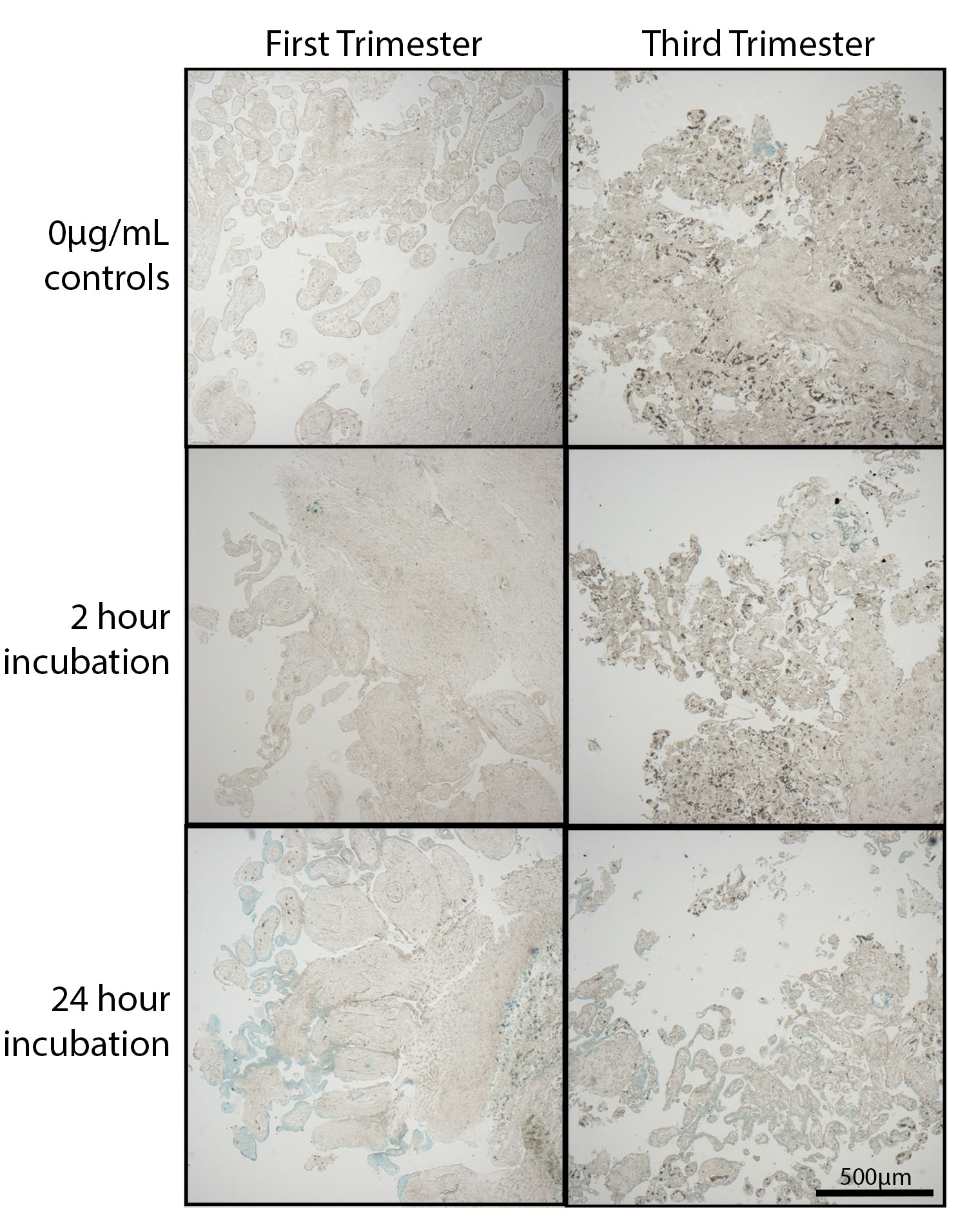

### Supplemental Figure 3

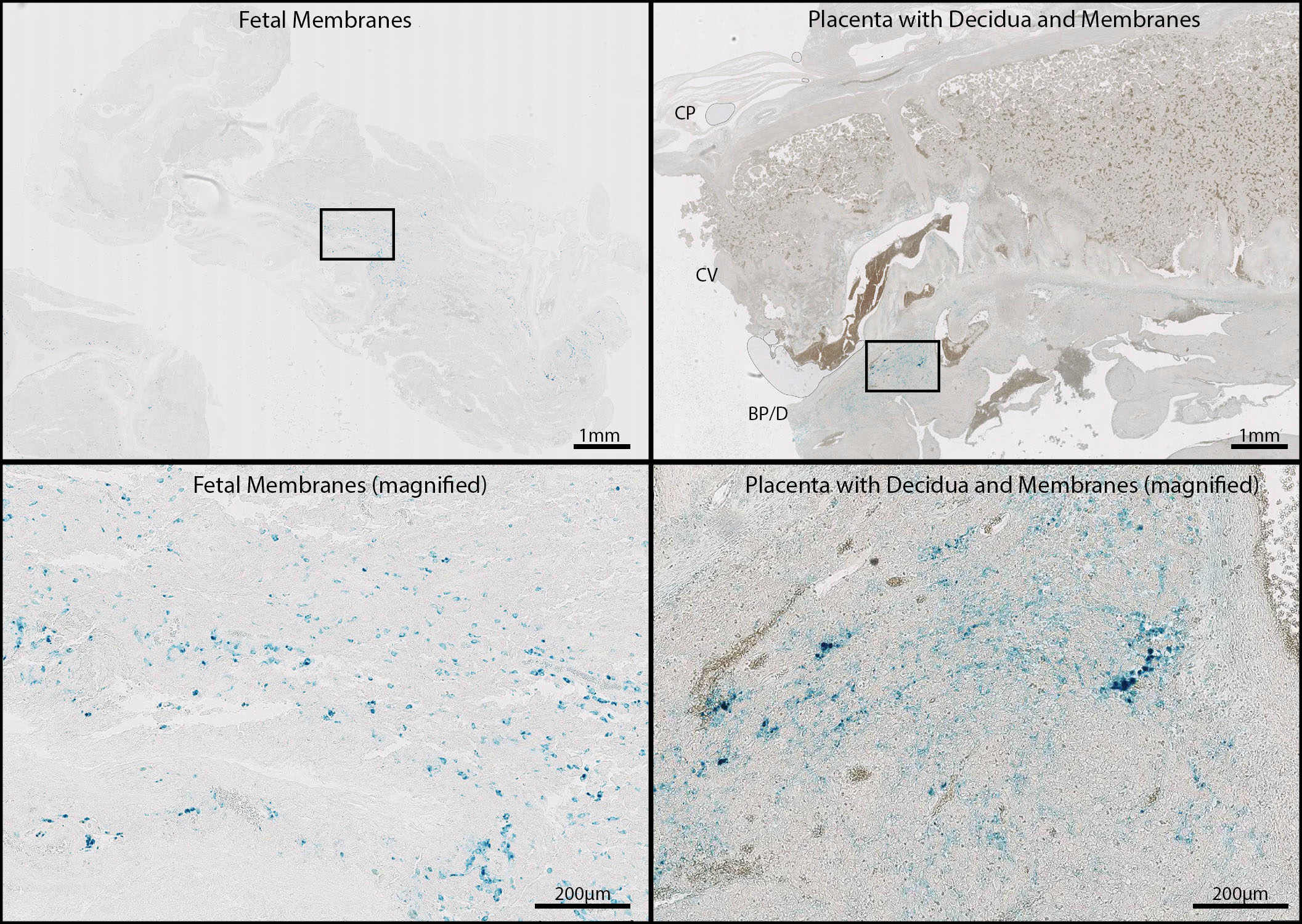
